# CD38 contributes to human natural killer cell responses through a role in immune synapse formation

**DOI:** 10.1101/349084

**Authors:** Mathieu Le Gars, Christof Seiler, Alexander W. Kay, Nicholas L. Bayless, Elsa Sola, Elina Starosvetsky, Lindsay Moore, Shai S. Shen-Orr, Natali Aziz, Purvesh Khatri, Cornelia L. Dekker, Gary E. Swan, Mark M. Davis, Susan Holmes, Catherine A. Blish

## Abstract

Natural killer (NK) cells use a diverse array of activating and inhibitory surface receptors to detect threats and provide an early line of defense against viral infections and cancer. Here, we demonstrate that the cell surface protein CD38 is a key human NK cell functional receptor through a role in immune synapse formation. CD38 expression marks a mature subset of human NK cells with a high functional capacity. NK cells expressing high levels of CD38 display enhanced killing and IFN-γ secretion in response to influenza virus-infected and tumor cells. Inhibition of CD38 enzymatic activity does not influence NK cell function, but blockade of CD38 and its ligand CD31 abrogates killing and IFN-γ expression in response to influenza-infected cells. Blockade of CD38 on NK cells similarly inhibits killing of tumor cells. CD38 localizes and accumulates at the immune synapse between NK cells and their targets, and blocking CD38 severely abrogates the ability of NK cells to form conjugates and immune synapses with target cells. Thus, CD38 plays a critical role in NK cell immune synapse formation. These findings open new avenues in immunotherapeutic development for cancer and infection by revealing a critical role for CD38 in NK cell function.

## Introduction

Natural killer (NK) cells are innate immune lymphocytes that are key players in the response to viral infections and contribute to tumor immune surveillance(Vivier et al., 2011). Due to their ability to kill tumor cells, NK cells are of great interest in cancer immunotherapy. NK cells express an array of inhibitory receptors, including the killer-cell immunoglobulin-like receptors (KIRs) and NKG2A, as well as activating receptors, including NKp46, NKp30 and NKG2D, which define NK cell maturation status and enable them to sense and respond to threats(Lee et al., 2007; Strauss-Albee et al., 2014; Vivier et al., 2008). NK cells can produce cytokines such as IFN-γ, which limit viral replication and tumor proliferation, and kill cells via release of cytolytic molecules or through engagement of death receptors(Vivier et al., 2008). In particular, the role and function of several receptors, such as NKG2D, NKG2A, and various KIRs have been studied in great detail in order to enhance NK cell cytotoxic responses to tumors(Baumeister et al., 2018; McWilliams et al., 2016).

CD38 is expressed on CD4 and CD8 T cells, monocytes, dendritic cells, and NK cells(Krejcik et al., 2016). In healthy human subjects, NK cell expression of CD38 is amongst the highest within peripheral blood mononuclear cells (PBMCs) (Ferrero and Malavasi, 1999; Krejcik et al., 2016; Malavasi et al., 1992). CD38 is often considered to be an activation marker on lymphocytes and possesses at least two intrinsic functions: 1) it is an ectoenzyme that transforms Nicotinamide Adenine Dinucleotide^+^ (NAD^+^) into cyclic Adenosyl-di-phosphate ribose (cADPR), and Nicotinamide Adenine Dinucleotide Phosphate (NADP) into Nicotinic Acid Adenine Dinucleotide Phosphate (NAADP), contributing to Ca^2+^ release from the endoplasmic reticulum(da Silva et al., 1998), and 2) it acts as an adhesion molecule, binding to CD31 and potentially other ligands (Deaglio et al., 1996, 1998; Horenstein et al., 1998; Nishina et al., 1994). CD38 adhesion to CD31 can promote extravasation of leukocytes to inflamed or infected tissues (Quarona et al., 2013).

Several lines of evidence suggest that CD38 plays a critical role in the activation of T and NK cells. Muñoz et al. demonstrated that CD38 is recruited to and accumulates in the immune synapse in Jurkat T-cell line in an antigen-specific and lck-dependent manner (Muñoz et al., 2008). In this T cell line, CD38 overexpression enhanced antigen-specific T cell activation, though it remained unclear whether the primary function of CD38 was in adhesion and stabilization of the immune synapse vs. localization of the enzymatic activity of CD38 to the immune synapse (Muñoz et al., 2008). CD38 ligation with agonistic antibodies induces calcium mobilization, cytokine secretion, and proliferation in a T cell receptor dependent manner, though differences exist between peripheral and tissue resident T cells in the specific signaling pathways used (Ausiello et al., 1996; Deaglio et al., 2001; Howard et al., 1993; Morra et al., 1998; Zubiaur et al., 1997, 1999). Further, there are suggestions that some CD38-mediated signaling functions may be independent of the T cell receptor and instead dependent on CD38 interacting with its ligand in some T cell lines (Cesano et al., 1998). In NK cells, Sconocchia et al. demonstrated that an ‘activating’ monoclonal antibody to CD38 redirects cytolysis and can induce NK cell degranulation and cytokine release in NK cells that had been pre-activated with IL-2 for 6 days (Sconocchia et al., 1999). Importantly, anti-CD38 antibody-mediated activation was not apparent in fresh NK cells, but only in those that had been pre-activated to induce expression of proteins required for cytolysis, and was independent of the intracellular calcium flux (Sconocchia et al., 1999). Subsequently, it was demonstrated that such antibody-mediated activation of NK cells required interaction with CD16 to mediate signaling (Mallone et al., 2001) and that CD38 and CD16 are functionally dependent and physically associated in human NK cells (Deaglio et al., 2002).

These data suggest that the function of CD38 on NK cells may be dependent on the manner in which CD38 is engaged and the context in which the NK cell is activated. Thus, it is intriguing to note that CD38 is upregulated on NK cells in the setting of viral infection and cytokine stimulation(Azeredo et al., 2006; Zidovec Lepej et al., 2003). Further, CD38 is part of a transcriptional signature highly diagnostic of human influenza infection(Andres-Terre et al., 2015). These observations led us to explore the function of CD38 in resting human NK cells responding to influenza-infected and tumor cells.

## Materials and methods

### Participants and study design

Six subjects within six days of the onset of laboratory-confirmed acute influenza A virus infection were recruited at the Stanford Emergency Department or Express Clinic as part of an approved protocol. Three individuals were infected by H1N1 virus strain and three with the H3N2 virus strain (determined by PCR). Blood was collected while patients were acutely infected, and a convalescent sample was collected at 28 days post infection. Healthy controls were obtained from de-identified LRS chambers purchased from the Stanford blood bank. Demographics are in Table S1. Phenotypic NK cell profiling was performed using two cohorts of healthy women recruited in separate years as part of a study aiming to characterize the impact of pregnancy on NK cell immune responses. The data from the non-pregnant healthy controls were used here. In the discovery cohort, thirty-two women were recruited at the Stanford Clinical and Translational Research Unit as part of influenza vaccine studies (NCT numbers: NCT01827462 and NCT03022422) between October 2012 and March 2013 (Table S2). In the validation cohort, twenty-one healthy women were studied between October 2013 and March 2014 from additional influenza vaccine studies (NCT numbers: NCT03020537, NCT03022422 and NCT02141581)(Table S3). For all functional assays, LRS chambers from healthy donors were purchased from the Blood Bank Center at Stanford University; both male and female subjects were used. For all subjects, PBMCs were isolated from whole blood by Ficoll-Paque (GE Healthcare) and cryopreserved in 90% fetal bovine serum (Thermo Scientific)/10% dimethyl sulfoxide (Sigma-Aldrich).

### PBMC Staining, CyTOF Acquisition, and analysis

Cryopreserved PBMCs from healthy individuals in both cohorts were thawed and cells were transferred to 96-well deep-well plates (Sigma), resuspended in 25 μM cisplatin (Enzo Life Sciences) for 1 min and quenched with 100% serum. Cells were stained for 30 min on ice, fixed (BD FACS Lyse), permeabilized (BD FACS Perm II), and stained with intracellular antibodies for 45 min on ice. Staining panels are described in Tables S4 and S5. All antibodies were conjugated using MaxPar X8 labeling kits (DVS Sciences). Cells were suspended overnight in iridium intercalator (DVS Sciences) in 2% paraformaldehyde in phosphate-buffered saline (PBS) and washed 1× in PBS and 2× in H_2_O immediately before acquisition on a CyTOF-1 (Fluidigm). Analysis of data was performed using FlowJo, version 9.9.4 (Tree Star).

### Virus Preparation and monocyte infection

A/California/7/2009 influenza (H1N1) wild-type influenza A virus was obtained from Dr. Kanta Subbarao, at the National Institutes of Health and propagated in embryonated chicken eggs as described(Karakus et al., 2018). Infection of monocytes was performed at multiplicity of infection (MOI) of 3 for 1 h at 37°C with 5% carbon dioxide as previously described (Kronstad et al., 2018).

### Fab preparation

Fab preparation was performed using the Mouse IgG1 Fab preparation micro kit (Thermo Scientific #44680) and CD38, CD31 and isotype control antibodies used in the functional experiments.

### Cell purification for functional assays

Cryopreserved PBMCs from healthy donors were thawed and washed with complete RP10 media (RPMI 1640 (Invitrogen) supplemented with 10% fetal bovine serum (FBS), 2 mM L-glutamine, 100 U/ml penicillin, 100 mg/ml streptomycin (Life Technologies)) and 50 U/mL benzonase (EMD Millipore). Autologous NK cells and/or monocytes were purified by magnetic-activated cell sorting via negative selection (Miltenyi). CD38^-^, CD38^+^ or bulk NK cells and/or autologous monocytes were sorted using Sony Sorter SH800 (Sony). The following antibodies were used to perform NK cell and monocyte sorting: CD3-Allophycocyanin (clone OKT3; BioLegend), CD14-Brilliant Violet 421 (clone HCD14; BioLegend), CD19-Alexa Fluor 488 (clone HIB19; Biolegend), CD38-PE (HIT2; Biolegend) and CD56-Phycoerythrin Cyanine 7 (clone NCAM; BioLegend).

### NK cell functional assays

NK cells were exposed to autologous H1N1-infected monocytes or K562 tumor cells at an effector:target (E:T) ratio 1:1 for 2 hours, followed by addition of 2 µM monensin, 3 µg/mL brefeldin A (eBiosciences), and anti-CD107a-allophycocyanin-H7 (BD Pharmingen) for 4 hours, followed by cell staining for flow cytometry analysis. When appropriate, NK cells were pre-incubated with activating IB4 antibody (IgG2a, gift from Dr. Fabio Malavasi, University of Torino) blocking CD38 (clone AT13/5, Bio-Rad), CD31 (clone WM59, Biolegend) and LFA-1 (clone TS1/22, Invitrogen) or isotype control antibodies or Fab.

### Cell staining and Flow-Cytometry Analysis

Cells were stained with LIVE/DEAD fixable Aqua Stain (Life Technologies), followed by surface staining and then fixed and permeabilized with FACS Lyse and FACS Perm II (BD Pharmingen) according to the manufacturer’s instructions. Cells were stained with anti-CD3-PE or -APC, anti-CD16-PerCPCy5.5 (clone 3G8; BioLegend), anti-CD38-PE or FITC (clone HIT2 Biolegend), anti-IFNγ-FITC or V450 (clone B27; BD Biosciences), anti-CD56-PEcy7, or anti-CD14-APC or -APC-H7 and fixed using 1% paraformaldehyde. Uncompensated data were collected using a three-laser MACSQuant® Analyser (Miltenyi). Analysis and compensation were performed using FlowJo flow-cytometric analysis software, version 9.9.4 (Tree Star).

### Flow cytometry-based conjugation assay

Isolated NK cells were labeled with CellTrace CFSE for 20 min at 37°C, washed and incubated with Fc Block for 5 min. NK cells were then incubated with no antibody, an isotype control, CD38, CD31 or LFA-1 blocking antibody for 15 min at 37°C. H1N1-infected monocytes or K562 cells were labelled with CellTrace Violet (CTV) for 20 min at RT. 10^5^ NK cells and 2 × 10^5^ target cells (E:T ratio 1:2) were mixed in 200 µl complete RP10, incubated for the indicated times (0 or 40 min) at 37°C, briefly vortexed, and fixed with 1% PFA in PBS. Cell mixtures were run on a three-laser MACSQuant® Analyser (Miltenyi). Analysis and compensation were performed using FlowJo flow-cytometric analysis software, version 9.9.4 (Tree Star). Percent of conjugated NK cells to their targets were calculated using this formula:

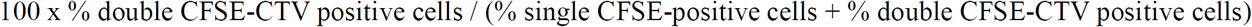

### Confocal microscopy

Isolated NK cells were stained using CellTrace Violet dye (Thermofischer) for 20 min at RT then washed twice in PBS. Isolated NK and mock- or H1N1-infected monocytes were incubated on a Poly-L-Lysine pre-coated 8 well µ-slide (Ibidi) for 2 hours. Cells were then washed in PBS-FBS 2%, fixed in PFA 4% for 15 min and washed twice in PBS-FBS 2%, then stained with mouse anti-CD38 and/or rabbit anti-LFA-1 antibody for 30 min at RT, then washed twice in PBS-FBS 2%. Secondary staining was performed using a goat anti-mouse AlexaFluor594 or a goat anti-rabbit AlexaFuor488 antibody for 30 min at RT. After washing the cells twice in PBS-FBS 2%, cell mount media (Ibidi) was added for microscopy. Images were acquired using LSM880 Meta (Zeiss) laser scanning confocal microscope equipped with a 63× (NA 1.4) DIC oil objective. All counting of synapses was performed in a blinded fashion.

### Statistical Analysis

Predictors of CD38^low^ vs. CD38^high^ NK cells were identified based on a Generalized Linear Model (GLM) with bootstrap resampling, in an immune cell specific implementation in R, as previously described in Kronstad et al., 2018 (Kronstad et al., 2018). Our open source R package *CytoGLMM* (Seiler et al., arxiv:1903.07976) is available here: https://christofseiler.github.io/CytoGLMM/.

Other statistical analyses were performed using GraphPad Prism, version 6.0d (GraphPad Software). The Wilcoxon matched pair test was used for all statistical analyses, except for the comparison between acutely-infected individuals and healthy controls which used the Mann-Whitney test.

## Results

### CD38 expression on NK cells is increased in response to influenza virus infection

Given that CD38 is highly expressed in NK cells (Fig. S1A), is a marker of lymphocyte activation, and is part of a highly accurate diagnostic of influenza infection, we hypothesized that it might be induced on NK cells during acute influenza infection. Indeed, acutely influenza-infected individuals displayed a very robust increase in CD38 expression on NK cells, which returned to baseline levels during convalescence (Fig. 1A, Table S1, Fig. S1B). As all of the healthy controls happened to be male, this raised the possibility that there may be sex-related differences in CD38 expression contributing to the differences observed. However, we observed no significant differences between men and women in NK cell expression of CD38, making this an unlikely possibility (Fig. S1C).

**Figure 1.**
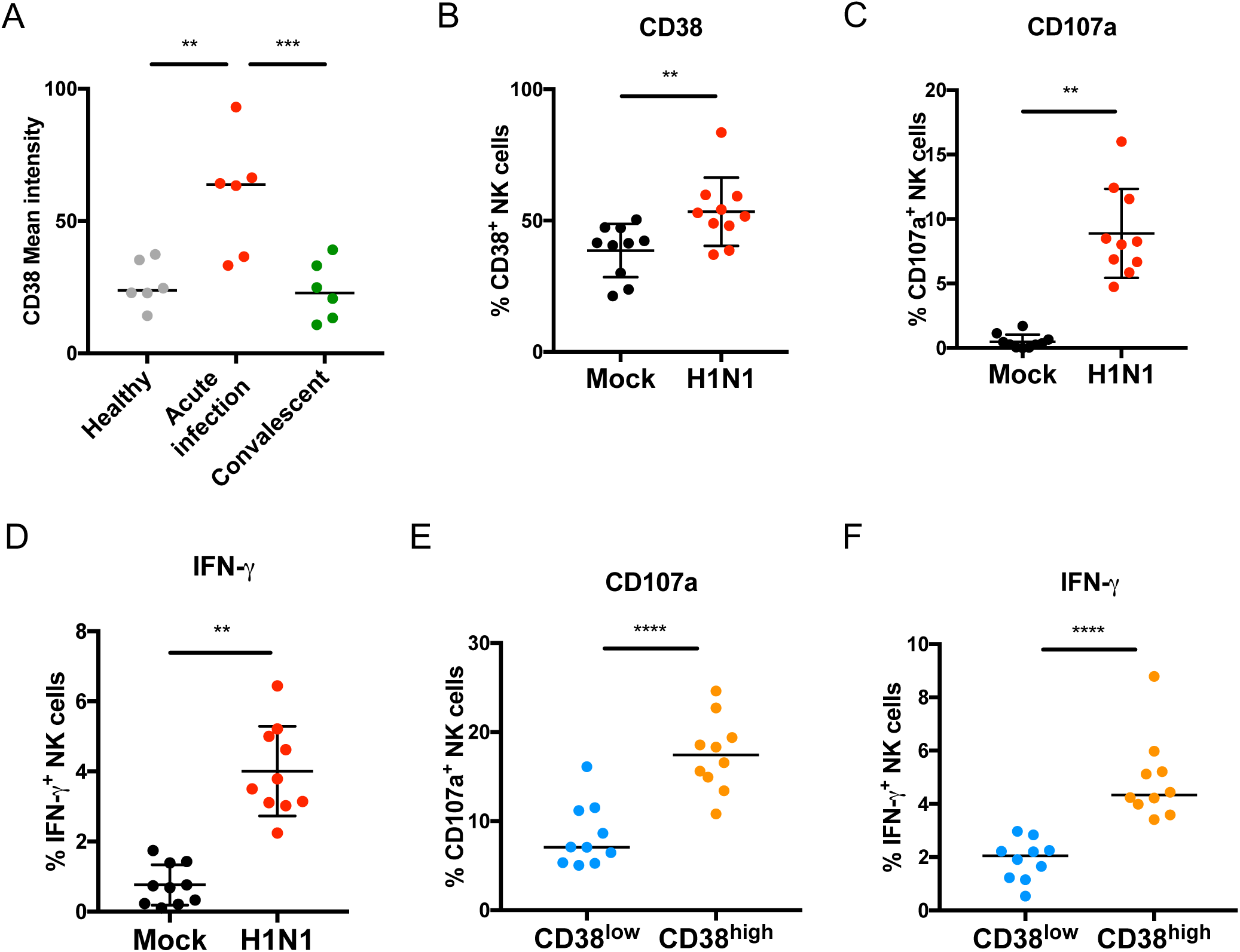
CD38 expression on NK cells is increased in response to influenza virus infection. (A) Median CD38 expression on NK cells from healthy donors, acutely influenza-infected donors, or the same influenza-infected donors in the convalescent phase. **P<0.01 (Mann-Whitney unpaired test to compare Controls vs. Acute infection) and ***P<0.001 (Wilcoxon matched-paired test to compare Acute infection vs. Convalescent). (B) CD38 expression on NK cells exposed to mock- or pandemic H1N1 influenza-infected monocytes. (C and D) CD107a and IFN-γ expression on NK cells from 10 healthy donors exposed to autologous H1N1-infected monocytes. **P<0.01. (E and F) CD107a and IFN-γ expression by CD38^-^ and CD38^+^ NK cells in response to influenza-infected monocytes. ****P<0.01.

To determine whether CD38 expression is similarly modulated on NK cells encountering infected cells *ex vivo*, we exposed NK cells to autologous influenza-infected monocytes as previously described(He et al., 2004; Kay et al., 2014; Kronstad et al., 2018). NK cells themselves are not infected by influenza in this system(Kronstad et al., 2018). We observed a significant increase in NK cell CD38 expression upon their encounter with monocytes infected with the pandemic A/H1N1 influenza strain (H1N1) compared to mock-infected conditions (Fig. 1B). We also observed significantly increased expression of CD107a, a marker of degranulation, and IFN-γ (Fig. 1C and D), demonstrating robust NK cell functional responses to the infected cells. The level of CD38 expression significantly correlated with CD107a induction (R^2^ = 0.79, p = 0.0006, Fig. S2A), though not significantly with IFN-γ expression (R^2^ = 0.30, p = 0.10, Fig. S2B). CD107a and IFN-γ expression were both significantly higher in CD38^high^ vs. CD38^low^ NK cells (Fig. 1E and F). Thus, CD38 expression is induced upon encounter with influenza-infected cells and its expression is associated with enhanced NK cell antiviral function.

### Profiling CD38^low^ and CD38^high^ NK cells

To understand how CD38 contributes to enhanced NK cell function, we explored the phenotype of CD38 expressing NK cells using mass cytometry (Fig. S1, Table S4). We used a Generalized Linear Model (GLM) with bootstrap resampling to identify markers predictive of CD38^high^ vs. CD38^low^ NK cells in a discovery cohort of 32 healthy women collected as part of a prior study(Kay et al., 2014; Kronstad et al., 2018; Le Gars et al., 2016). CD38^high^ NK cells were more likely to express several activating (CD11b, CD244, HLA-DR, NKp30, NKp44 and NKp46) and inhibitory (KIR2DL1, KIR2DL3, KIR2DL5, KIR3DL2 and LILRB1) receptors, while CD38^low^ NK cells were more likely to express the activating receptor NKG2D, and the inhibitory receptors NKG2A and KIR2DL4 (Fig. 2A). A similar analysis of 21 healthy women in a validation cohort using a different antibody panel confirmed the increased expression of NKp30 and NKp46, but not NKp44, in CD38^high^ NK cells compared to CD38^low^ NK cells (Fig. 2B, Table S4, Table S5). We also confirmed the decreased expression of NKG2A, but not NKG2D, among CD38^high^ NK cells in this validation cohort. Expression of perforin, which was only evaluated in the validation cohort, was significantly predictive of CD38^high^ NK cells. Manual gating confirmed these results for both cohorts (Fig. S3, S4, S5A and S5B). Overall, the increased expression of several KIRs, NKp30, NKp46, CD11b, and perforin, along with decreased expression of NKG2A, suggest that CD38^high^ NK cells are a mature subset with high cytotoxic potential.

**Figure 2.**
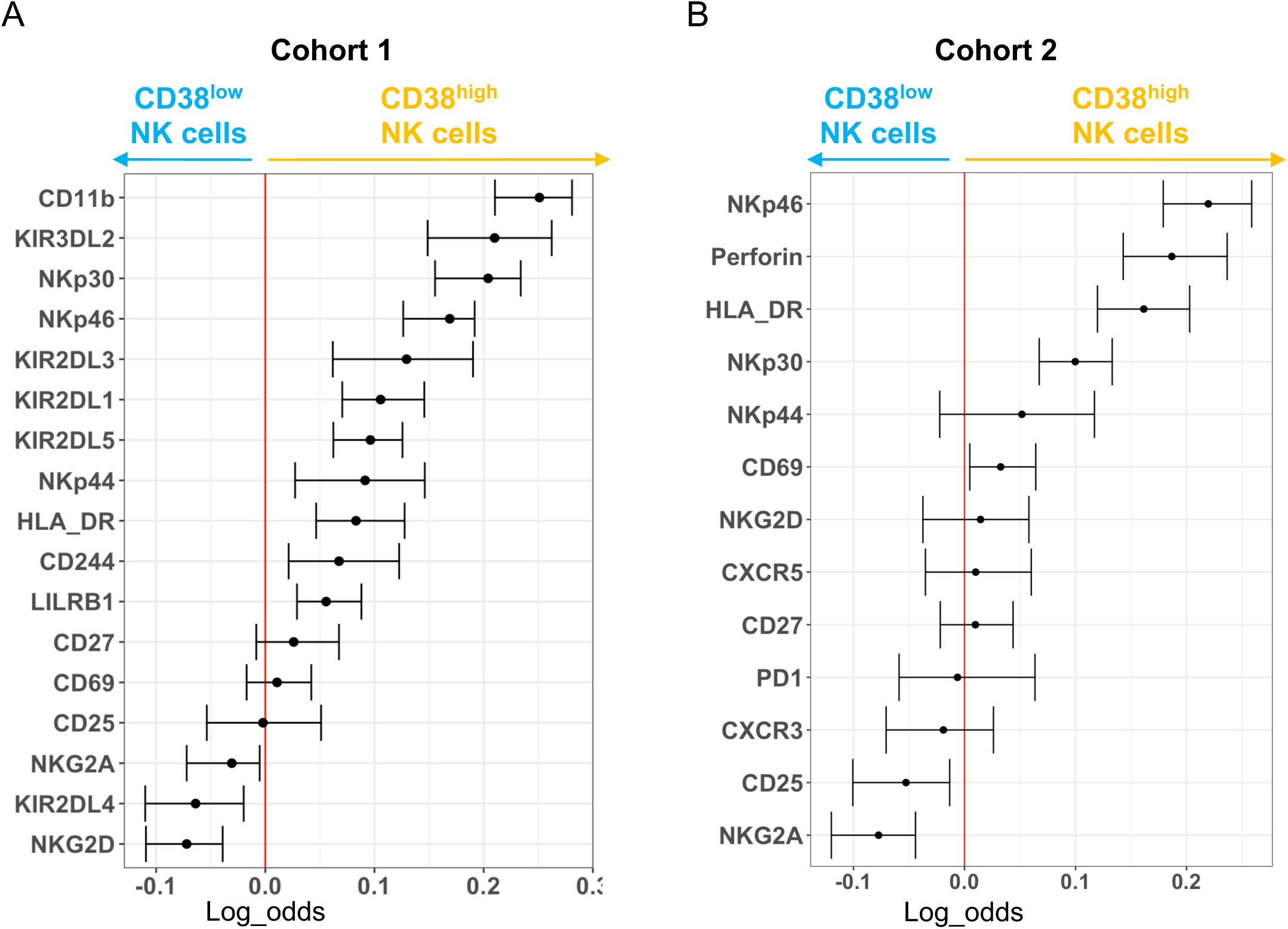
Deep profiling of CD38^low^ and CD38^high^ NK cells from healthy individuals. A GLM with bootstrap resampling was used to identify markers predictive of CD38^low^ vs CD38^high^ NK cells within the discovery cohort (A) and validation cohort (B). The markers studied are listed vertically and the x-axis represents the log-odds that expression of that marker predicts the given subset.

### CD38 plays a direct role in enhanced NK cell function to influenza-infected cells

To determine whether CD38-expressing NK cells have any enhanced functions, we sorted CD38^low^ and CD38^high^ NK cells and assessed a number of immune response indicators (Fig. S6A). We found that compared to CD38^low^ cells, CD38^high^ NK cells had a 3.5-fold greater expression of CD107a, 7.2-fold greater IFN-γ expression, and 2.2-fold greater ability to induce the death of influenza infected monocytes (Fig. 3A, B and C). To directly test the role of CD38 in this enhanced function, we assessed the potential contributions of CD38-CD31 mediated adhesion using the previously reported agonistic anti-CD38 antibody IB4 and blocking antibodies to both CD38 and its ligand, CD31, which is expressed on monocytes (Fig. S6B). Given the critical role of LFA-1 in immune synapse formation(Orange, 2008), blocking antibodies to LFA-1 were also used as a positive control. In a distinction from the setting of IL-2 activated NK cells, the agonistic IB4 antibody did not have a significant effect on NK cell function in response to influenza-infected cells (Fig. S6C). However, blocking CD38 and its ligand CD31 led to a marked reduction in NK cell function in response to influenza-infected monocytes, with 2.4-fold reduction in CD107a expression, 2.2-fold reduction in IFN-γ production, and 1.7-fold reduction in killing of infected monocytes with CD38 blocking (Fig. 3D, E and F). Blocking CD31 and LFA-1 also significantly abrogated NK cell expression of CD107a and IFN-γ. In light of concerns for blocking antibodies driving antibody-dependent cellular cytotoxicity via the FcRγIII receptor (CD16) on NK cells, we repeated the experiments with Fabs derived from CD38 and CD31 blocking antibodies. Both Fabs significantly decreased the NK cell CD107a and IFN-γ response to influenza-infected monocytes, just as the parent blocking antibodies did (Fig. S6D and E). We next explored whether the enzymatic activity of CD38 might play a role in NK cell function to influenza-infected cells. The CD38 enzymatic inhibitor, kuromanin, did not significantly alter NK cell function at concentrations known to inhibit this function in T cells(Morandi et al., 2015)(Fig. S6F and G). Thus, CD38 appears to contribute to NK cell responses against infected cells through its role in binding to CD31.

**Figure 3.**
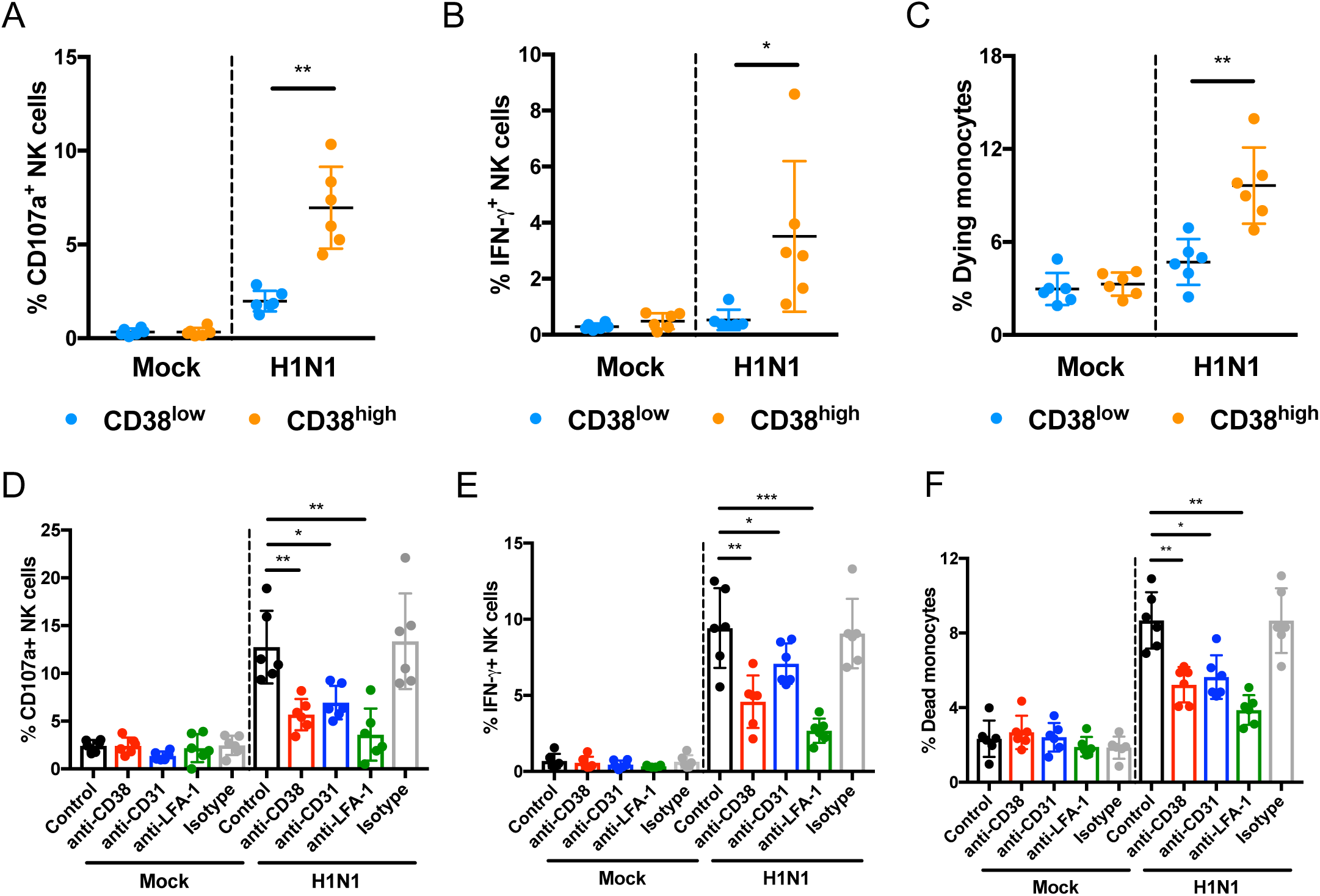
CD38 plays a direct role in NK cell immune response to influenza infection. NK cell expression of CD107a (A), IFN-γ (B), and the frequency of dying monocytes based on a viability dye (C) from co-cultures between sorted populations of CD38^low^ and CD38^high^ NK cells and autologous monocytes infected with H1N1 influenza-virus or mock-infected (N=6 healthy blood bank donors). *P<0.05, **P<0.01. (D to F) Isolated NK cells from healthy donors (N=6) were exposed to autologous H1N1-infected monocytes in presence or absence of an isotype control antibody or a CD38, CD31 or LFA-1 blocking antibody or no antibody. CD107a expression (D), IFN-γ production (E) and monocyte killing (F) by NK cells were measured by flow cytometry. *P<0.05, **P<0.01, ***P<0.001.

### CD38 is necessary for the establishment of an immune synapse with influenza-infected cells

CD38-CD31 interactions promote adhesion and rolling of leukocytes to endothelial cells during extravasation(Deaglio et al., 2000) and may play a role in synapse formation by T cells (Muñoz et al., 2008). This led us to investigate whether CD38-CD31 interactions play a role in the formation of the immune synapse between NK cells and infected cells. Blocking antibodies to both CD38 and LFA-1, which was used as a positive control, led to a significant 1.89- and 1.97-fold reduction in the formation of NK cell-monocyte conjugates compared to untreated cells or isotype control antibody-treated cells, respectively (Fig. 4A and Fig. S7A), suggesting that CD38 is important for adhesion between NK cells and their targets. Using confocal microscopy, we found that CD38 localizes to the immune synapse at the contact zone between NK cells and H1N1-infected monocytes, which was identified based on co-staining with the adhesion molecule LFA-1 (Fig. 4B and C). Minimal accumulation of CD38 was observed when NK cells were in contact with mock-infected monocytes. In the presence of blocking CD38 or CD31 antibodies, the formation of the immune synapse was strongly and significantly inhibited compared to cells incubated with an isotype control antibody (2.7- and 2.0-fold reduction, respectively, Fig. 4D and E). Similarly, immune synapse formation was rarely observed between H1N1-infected cells and CD38^low^ NK cells. Together, these data indicate that CD38 contributes to the establishment of the immune synapse between NK cells and influenza-infected cells, which is critical for NK cell immune responses.

**Figure 4.**
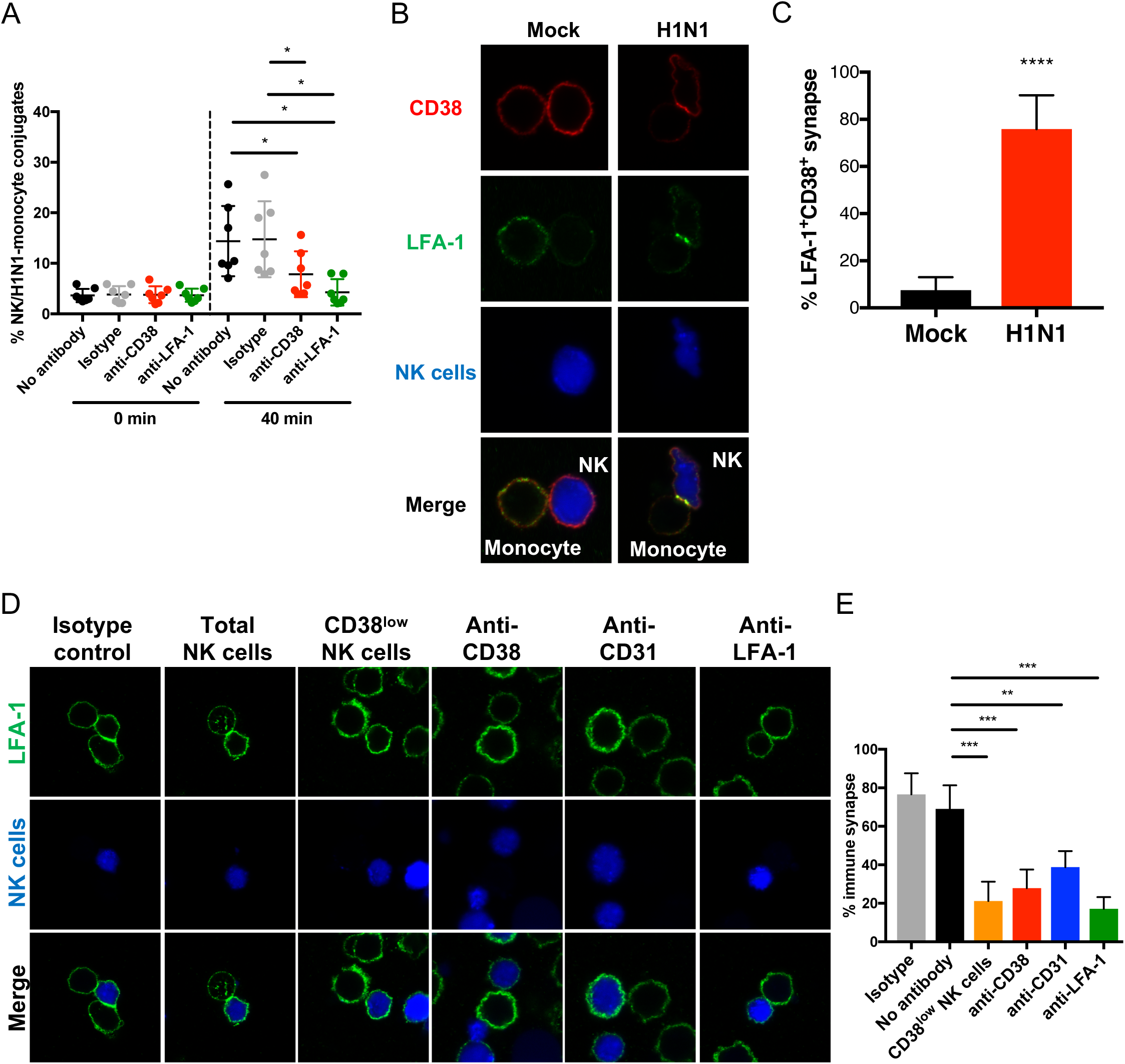
CD38 contributes to the immune synapse between NK cells and H1N1-infected cells. (A) NK cell:monocyte conjugate formation was assessed by flow cytometry in the presence of no antibody, isotype control antibody, or blocking antibodies to CD38 and LFA-1. N=6, *P<0.05. (B and C) Immune synapse formation was determined by the accumulation of LFA-1 at the contact zone (N=30) between NK cells (in blue) and mock- or H1N1-infected monocytes, and CD38 co-localization at the immune synapse. (B) The image depicted is representative of the accumulation of CD38 at the immune synapse (LFA-1 colocalization). (C) The percentage of immune synapses with co-localization is represented based on blinded counting. ****P<0.0001 (Wilcoxon matched-paired test). (D and E) Quantification of the frequency of immune synapse formation between total NK cells and influenza-infected monocytes in the presence of blocking antibodies or sorted CD38^low^ NK cells. 60 contacts between NK cells and H1N1-infected monocytes were observed and LFA-1 accumulation was used to identify immune synapse formation in a blinded fashion. (D) A representative image is depicted for each condition. (E) The percentage of immune synapse for each condition is represented. **P<0.01 and ***P<0.001.

### CD38 contributes to immune synapse formation between NK cells and cancer cells

To explore whether the role of CD38 extends beyond the response to influenza-infected cells, we examined NK cell responses to K562 tumor cells (Lisovsky et al., 2015). NK cells upregulated CD38 in response to K562 cells (Fig. S8A). Sorted CD38^high^ NK cells were significantly better than CD38^low^ NK cells at killing tumor cells (Fig. 5A and B) and secreting IFN-γ (Fig 5C). Blocking CD38 significantly diminished CD107a expression on NK cells and inhibited tumor cell killing (Fig. 5D and E), but not IFN-γ production (Fig. 5F). Blocking CD31 had no significant effect on CD107a expression, K562 cell death, or IFN-γ production, suggesting that CD38 may bind an alternate ligand on these tumor cells. Blocking CD38 or LFA-1 significantly diminished conjugate formation with tumor cells (Fig. 5G and Fig. S8B). Further, sorted CD38^high^ NK cells have a significantly greater potential to form conjugates with K562 cells than do CD38^low^ NK cells (Fig. 5H and Fig. S8C). As in the synapse with infected cells, CD38 polarizes to the immune synapses between NK cells and K562 tumor cells (Fig. 5I and Fig S8D). These results indicate that CD38 plays a crucial role in the establishment NK cell cytotoxic function and immune synapse formation with cancer cells.

**Figure 5.**
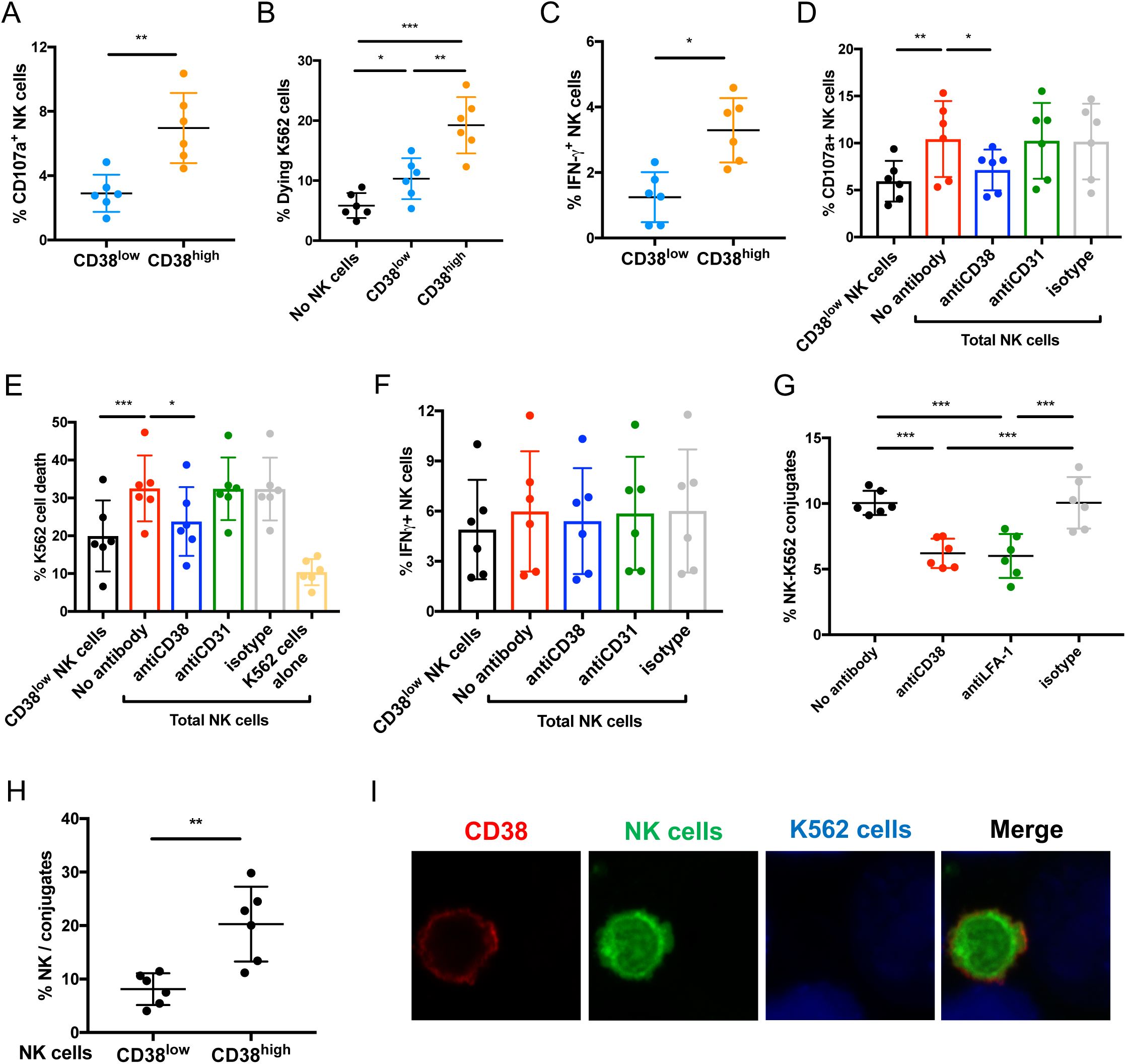
CD38 contributes to immune synapse formation between NK cells and cancer cells. Functional responses of CD38^low^ and CD38^high^ NK cells to K562 tumor cells assessed by the frequency of CD107a expression (A), the frequency of dying monocytes (B), and the frequency of IFN-γ producing NK cells (C). N=6 healthy blood bank donors. *P<0.05, **P<0.01 and ***P<0.001. Functional response of NK cells to K562 tumor cells based on CD107a expression (D), the frequency of dying tumor cells (E), and the IFN-γ production (F) within sorted CD38^low^ NK cells or total NK cell populations treated or not with isotype control antibody or blocking antibodies to CD38 and CD31. N=6 healthy donors. *P<0.05, **P<0.01, ***P<0.001. (G) NK cell-K562 cell conjugation was determined by flow cytometry in the presence of no antibody, isotype control antibody, or blocking antibodies to CD38 and LFA-1. N=6 healthy donors. ***P<0.001. (H) CD38^low^ and CD38^high^ NK cells were sorted from healthy blood donors (N=6) and stained with CFSE. In parallel, K562 cells were stained with CellTrace Violet. NK cells were then exposed to K562 cells for 40 min. NK cell-K562 cell conjugation was determined by the accumulation of the double positive CFSE^+^CellTrace Violet^+^ population by flow cytometry. Graph represents a summary of the experiments. **P<0.01 (Wilcoxon matched-paired test). (I) Confocal imaging of NK cells with K562 tumor cells. NK cells are stained with CFSE, K562 cells with CellTrace Violet, and CD38 with AlexaFluor594. A representative image demonstrating CD38 polarization to the immune synapse is depicted.

## Discussion

The discovery and characterization of new receptors involved in NK cell function has the potential to improve our efforts to develop therapeutics for viral infection and cancer. Here, we demonstrate a role for the cell surface protein CD38 in human NK cell function. Prior studies revealed that antibody-mediated cross-linking of CD38 on IL-2 stimulated NK cells lead enhanced NK cell degranulation and activation, yet such cross-linking had no effect on resting human NK cells (Sconocchia et al., 1999)(Mallone et al., 2001)(Deaglio et al., 2002). Here, using mass cytometry and functional assessments, we demonstrate that CD38 expression marks a mature subset of NK cells with high cytolytic and cytokine producing capacity. Further, our study reveals that CD38 plays a critical role in NK cell function against both viral-infected and tumor cells by enhancing immune synapse formation. Such activity is present in resting human NK cells directly ex vivo and does not require prior activation. These data reveal a novel mechanism regulating human NK cell immune responses.

CD38 is commonly viewed as a lymphocyte activation marker; its increased expression on NK cells responding to infected and tumor cells could merely be a consequence of NK cell activation. In fact, CD38 expression signifies a population of NK cells with increased expression of KIRs, CD11b, CD57, and perforin, suggesting a high degree of maturation(Luetke-Eversloh et al., 2013). In particular, the high expression of KIRs and perforin suggests that CD38^high^ NK cells are highly functionally competent with high cytotoxic capacity(Anfossi et al., 2006; Elliott and Yokoyama, 2011). Prior studies suggested that CD38 may contribute to NK cell activation: cross-linking CD38 on IL-2 activated NK cells promoted NK cell activation and degranulation in a CD16-dependent manner (Sconocchia et al., 1999)(Mallone et al., 2001)(Deaglio et al., 2002). It is important to note that in the prior studies, cross-linking CD38 on resting NK cells failed to activate NK cells, and it remains possible that it was not CD38 cross-linking, but rather NK cell activation to kill other NK cells via antibody-dependent cellular cytotoxicity against anti-CD38 antibody-coated NK cells (fratricide).

Here we identify a distinct role for CD38 on resting NK cells that is suggested by data indicating that CD38-CD31 interactions may contribute to T cell immune synapse formation (Muñoz et al., 2008). In the setting of the response to influenza-infected cells, cross-linking CD38 with agonistic antibodies did not significantly alter the response, nor did inhibiting the enzymatic activity of CD38. Instead, our data show that CD38 contributes to the formation of immune synapse between NK cells and their targets, even on freshly isolated human NK cells in the absence of prior stimulation, revealing a novel role for this molecule on NK cells. CD38-CD31 interactions are necessary to establish the NK cell immune synapse and for full cytotoxic activity against influenza-infected cells. Importantly, CD38 plays a similar role in the NK cell response to K562 tumor cells, but it appears that other ligands might also be involved, as blocking CD31 did not significantly inhibit K562 killing. CD38 has been postulated to have other ligands such as CD9(Zilber et al., 2005), and identification of such ligands will be an area of future investigation. Interestingly, the IFN-γ response to tumor cells, unlike killing activity, was not dependent on CD38 expression. This may relate to the local cytokine environment, as influenza-infected monocytes produce abundant IFN-α that drives NK cell IFN-γ secretion(Kronstad et al., 2018), but K562 cells do not produce such IFN-α. Together, these data indicate that CD38 has a previously unrecognized but critical role in facilitating NK cell cytolytic activity against both infected cells and tumor targets.

NK cell immune synapse formation occurs in three successive stages: recognition, effector, and termination(Orange, 2008). During the recognition phase, CD2 and LFA-1 interactions with CD58 and ICAM, respectively, are critical steps in the early phases of immune synapse formation that aid in adhesion (Krensky et al., 1983)(Comerci et al., 2012; Orange, 2008)(Altomonte et al., 1993)(Altomonte et al., 1993; Orange, 2008; Wang et al., 2015). We propose that CD38-CD31 interactions are an additional component of NK cell immune synapse formation, as had been previously suggested in T cells (Muñoz et al., 2008). Identifying the exact stage in which CD38 interacts with CD31 or other ligands during immune synapse formation will be a critical step forward in our understanding of the precise mechanisms. CD2, LFA-1 and CD38 may also physically interact and serve to stabilize or enhance NK cell adhesion to their targets. Defining the nature of such interactions will be an important area of further study.

During the effector phase, the immune synapse plays an essential role in NK cell activation. This activation is thought to be partly mediated by LFA-1 intracellular signaling(Barber et al., 2004; Orange, 2008). CD38 has a short cytoplasmic domain without any known signaling motifs, but can potentiate signaling through the CD3-TCR complex in T cells, MHC class II in monocytes, CD19 in B cells(Lund et al., 1996; Zilber et al., 2005; Zubiaur et al., 2002), and CD16 in NK cells (Sconocchia et al., 1999)(Mallone et al., 2001)(Deaglio et al., 2002). The co-localization of CD38 with LFA-1 in the immune synapse suggests that CD38 could lead to synergistic effects on LFA-1 signaling in NK cells. Consistent with this idea, LFA-1 signaling is in part mediated through the phosphorylation of LCK and ZAP-70, two adaptor proteins that are phosphorylated by the interaction of CD38 and the CD3-ζ subunit in T cells(Muñoz et al., 2003; Verma and Kelleher, 2017). Thus, CD38 could both strengthen adhesion and enhance LFA-1 signaling, explaining the importance of both proteins in the formation of the immune synapse.

Harnessing and directing the cytotoxic activity of NK cells has been of great interest within the scientific community over the last several years. Several studies suggest that NK cells with antibody specificities may be as effective as CAR-T cells in recognizing and killing tumor cells (Liu et al., 2017; Rezvani et al., 2017). Thus, the identification of a new role for CD38 in immune synapse formation and optimal cytotoxic activity has significant implications for these therapies. It will be important to understand the factors that control CD38 expression *in vivo* to assure its retention on engineered cells. For instance, we find that both IL-2 and IL-15, two important cytokines for NK cell homeostasis, promote increased CD38 expression *in vitro* (data not shown). These data also raise interesting questions about the CD38-targeting agents that are currently in clinical use, primarily for multiple myeloma and chronic lymphocytic leukemia(Damle et al., 1999; Lokhorst et al., 2015). Clinical treatment with daratumumab, a specific human CD38 binding antibody, improves patient outcomes(Lokhorst et al., 2015), but also may lead to NK cell fratricide(Frerichs et al., 2018; Wang et al., 2018). It is currently unclear how daratumumab affects CD38-mediated immune synapse formation, and how this influences NK cell killing. Further, several forms of human glioblastoma have been shown to overexpress CD31(Musumeci et al., 2015), potentially making them excellent targets for NK cells with enhanced CD38 expression.

A limitation of our study is the fact that our mass cytometry panels differed between the two cohorts and that this phenotypic data was exclusively on female subjects. To ameliorate this concern, all functional experiments were performed with samples from healthy blood bank donors of both sexes. A second limitation is that we used blocking antibodies or Fab and sorted populations rather than knockouts to establish CD38 function. While CRISPR-Cas9-mediated editing of NK cells is feasible(Phan et al., 2016; Somanchi and Lee, 2016), the expansion protocol required leaves NK cells unable to respond to influenza-infected cells (not shown). Thus, even if we were able to knockout CD38, we could not appropriately assess its impact on NK cell function in this physiologic antiviral response.

In summary, our goal was to better understand the role of CD38 in NK cell biology and activity in the context of influenza A virus infection and cancer. We find that, at least *in vitro*, CD38 expression is crucial in both settings. We also demonstrate a novel role for CD38 in the establishment of immune synapses that are characteristic of efficient killing by NK cells. This will be useful for potential NK cell engineering and other therapeutic applications.

## Acknowledgements

We thank our study volunteers for their participation, Sally Mackey for regulatory and data management, Sue Swope for consenting and conducting study visits, and the staff of the Stanford Vaccine Program for overall study coordination.

## Author contributions

M.L.G., A.W.K., N.L.B. and C.A.B. designed experiments. M.L.G., A.W.K. and N.L.B. E. Sola. conducted experiments. M.L.G. and C.S. analyzed the data. L.M., S.S.S.O., P.K., collaborated and provided advice in the analysis of the data. M.M.D., C.L.D., G.E.S. and N.A. coordinated the clinical studies and provided human samples. M.L.G and C.A.B. wrote the manuscript; all authors contributed revisions and edits.

