## Supplementary figures and images for "CD38 contributes to human natural killer cell responses through a role in immune synapse formation"

### Supplemental_figures

#### Figure S1

A

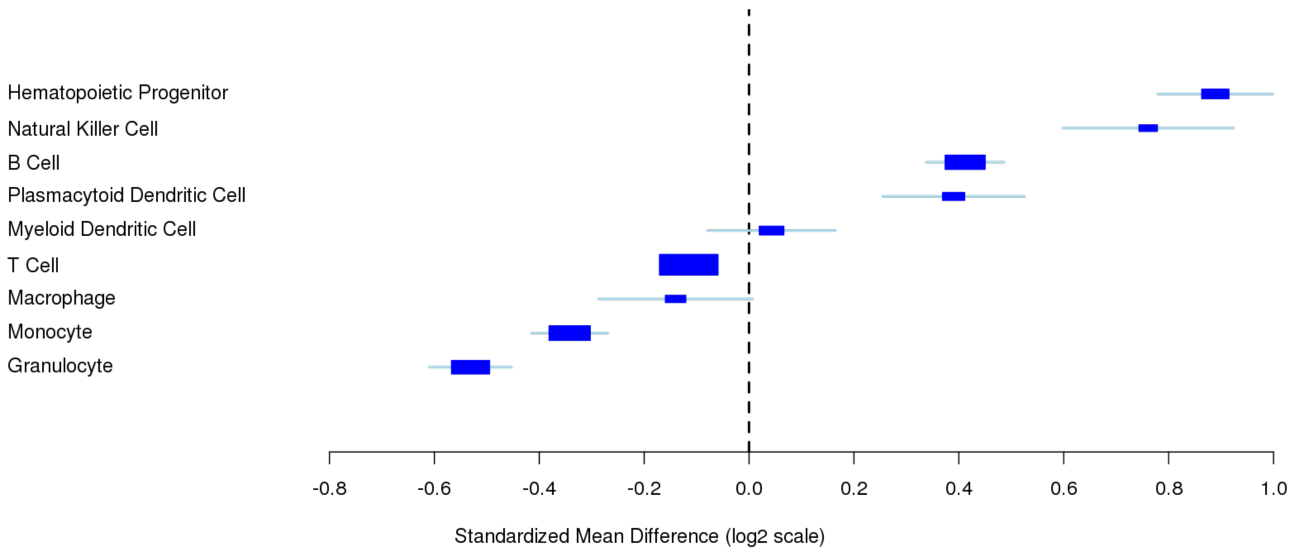


B

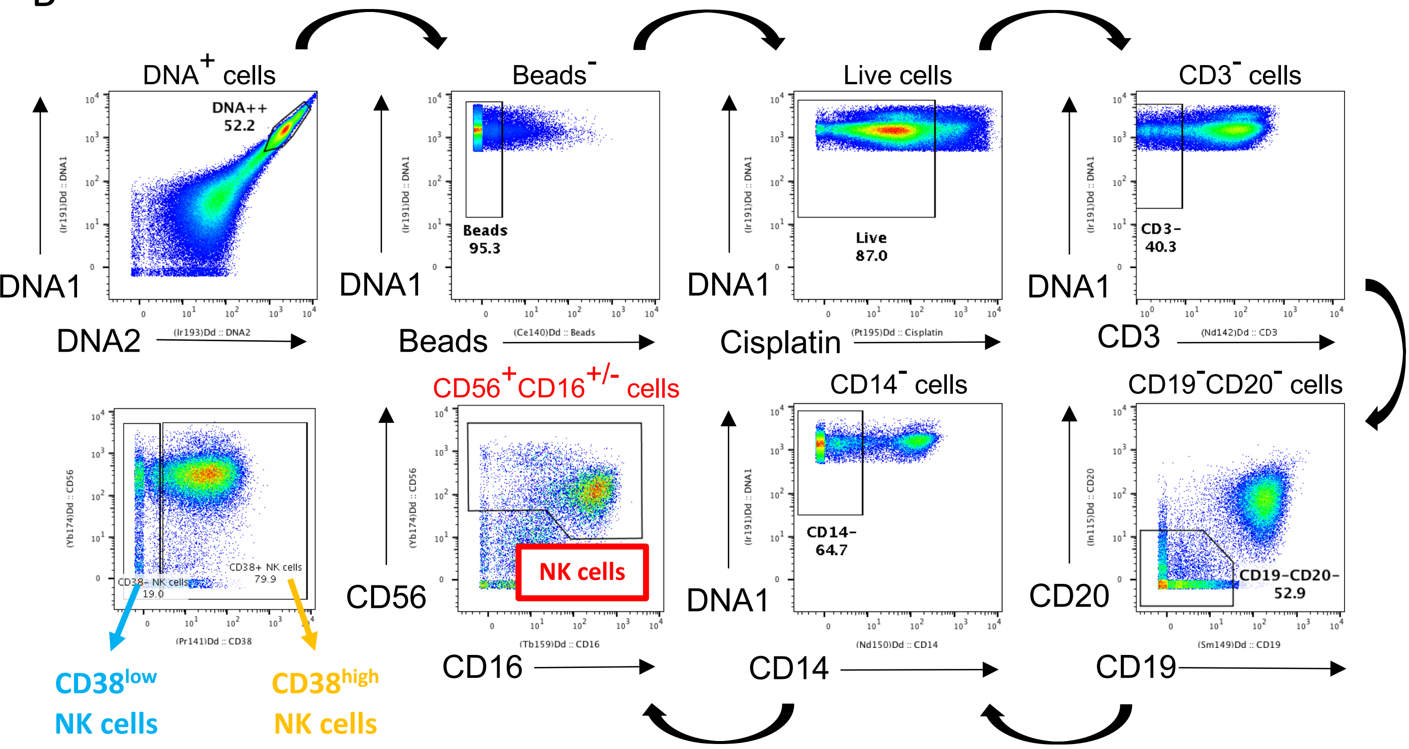


C

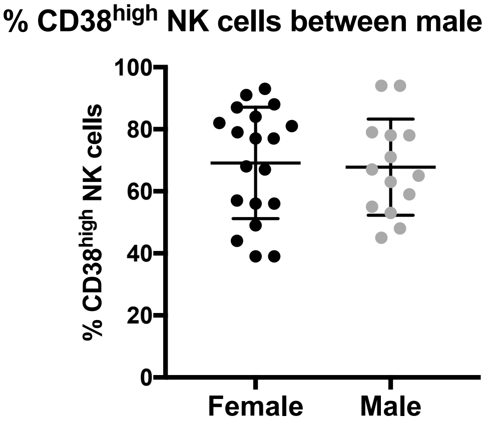

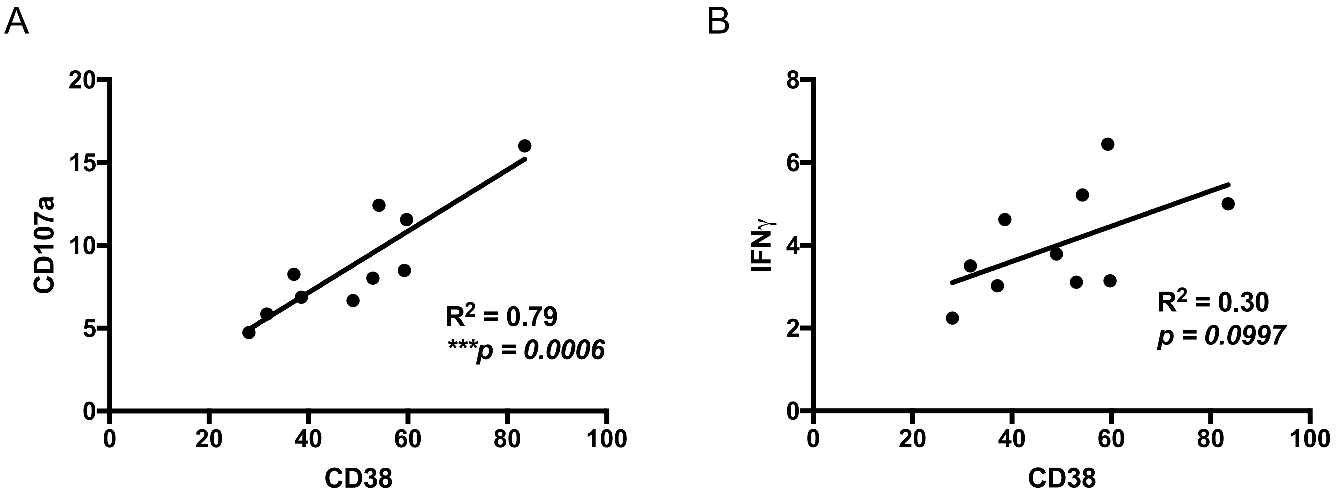

#### Figure S3

##### Cohort 1

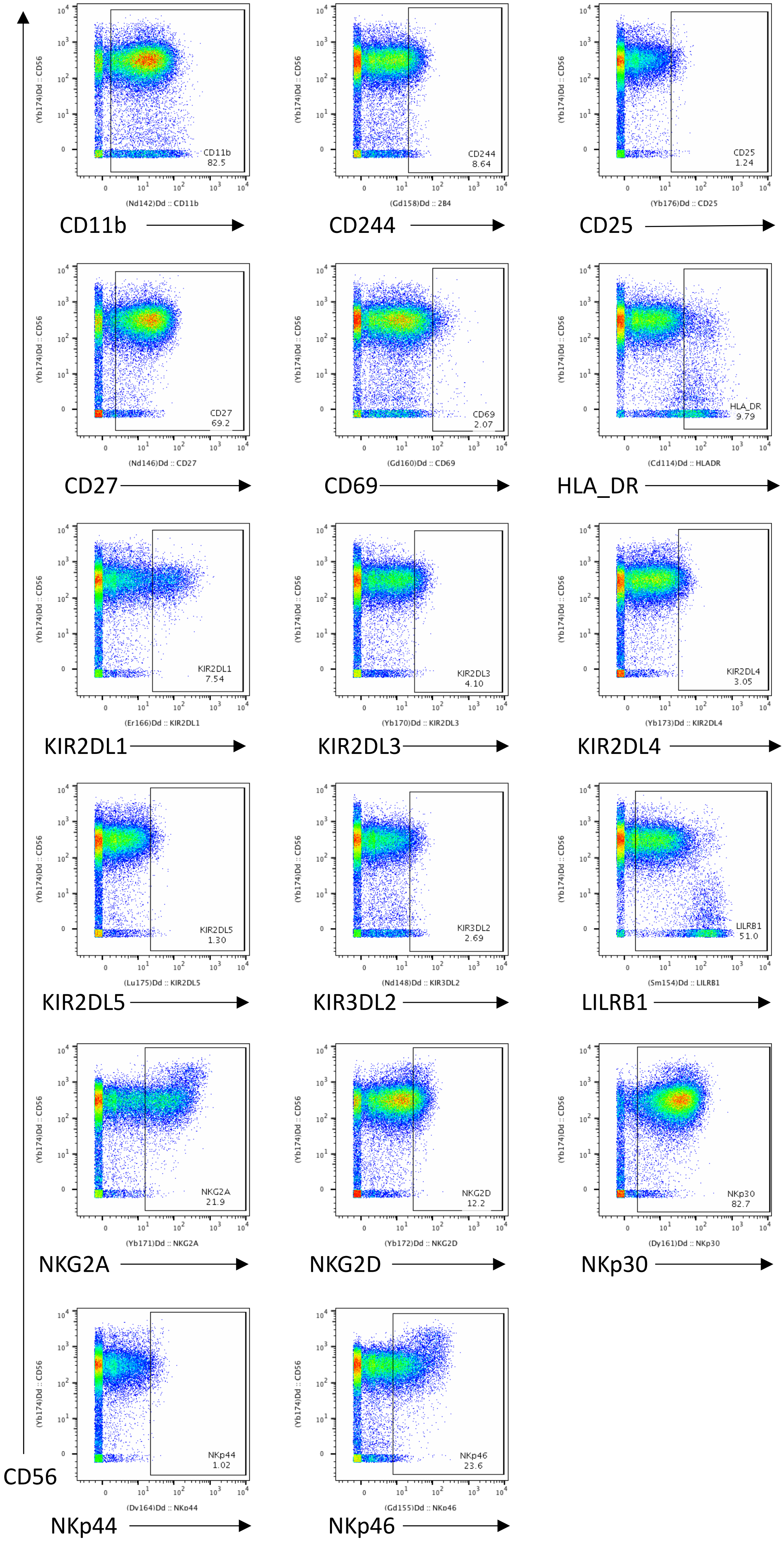

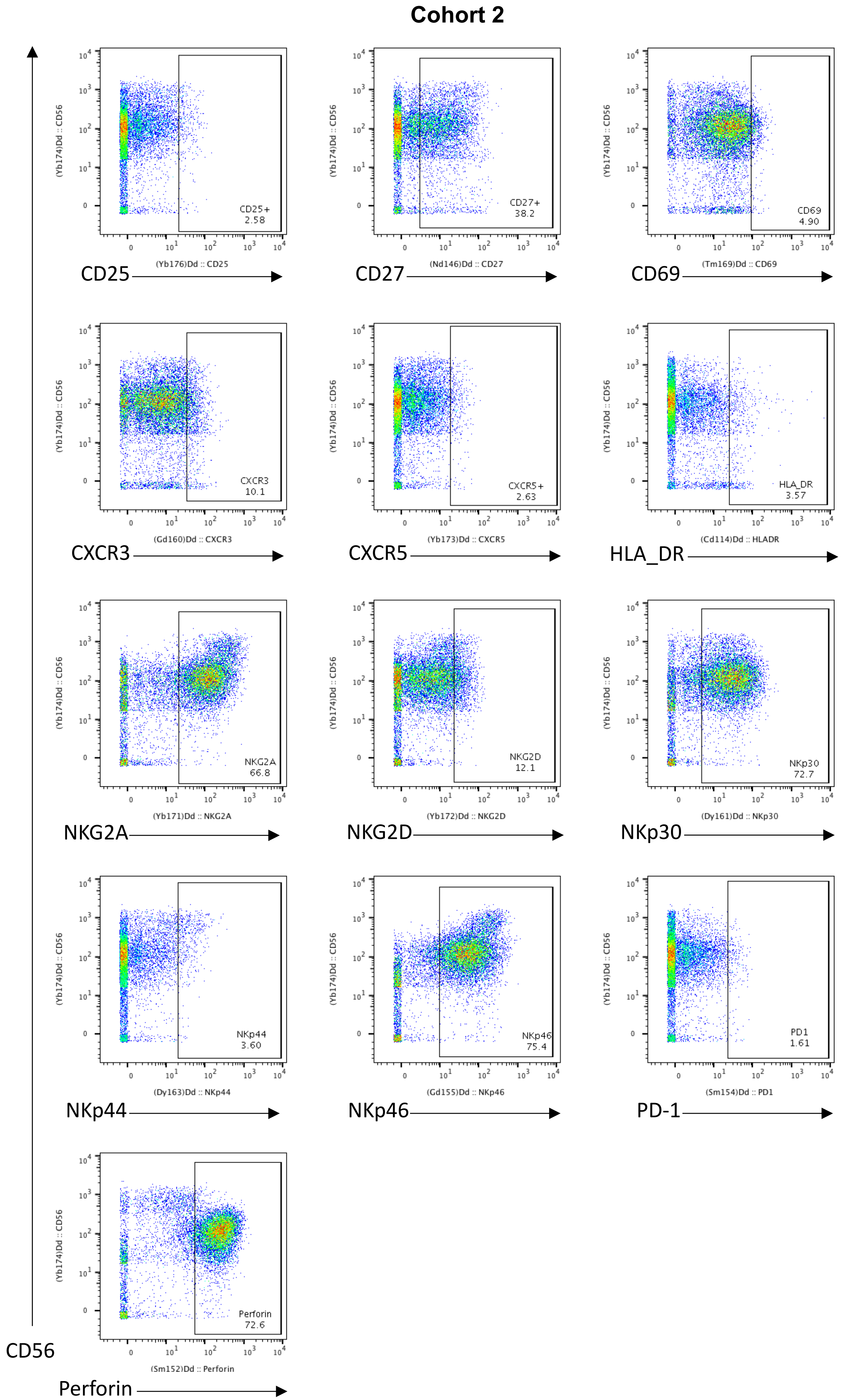

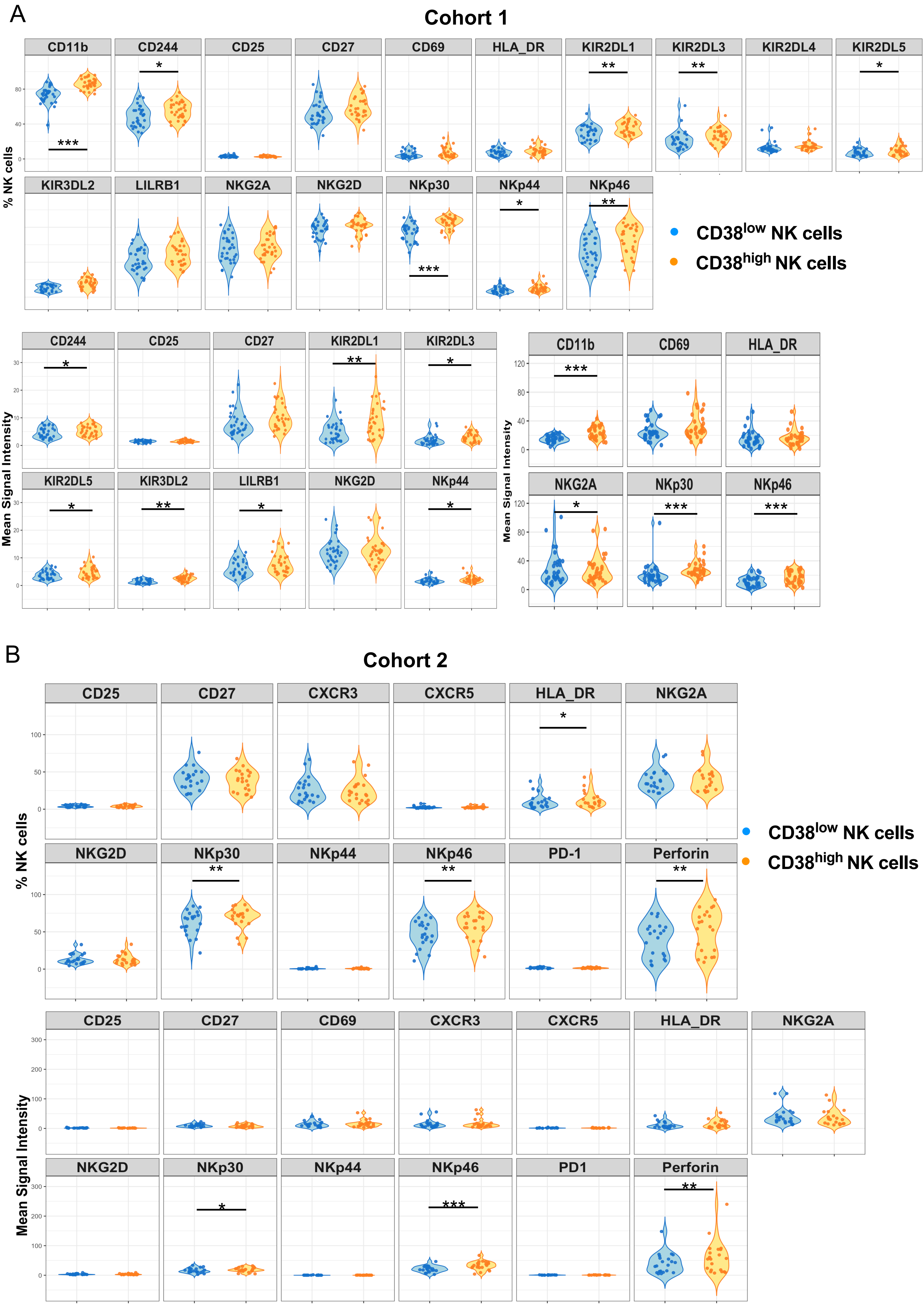

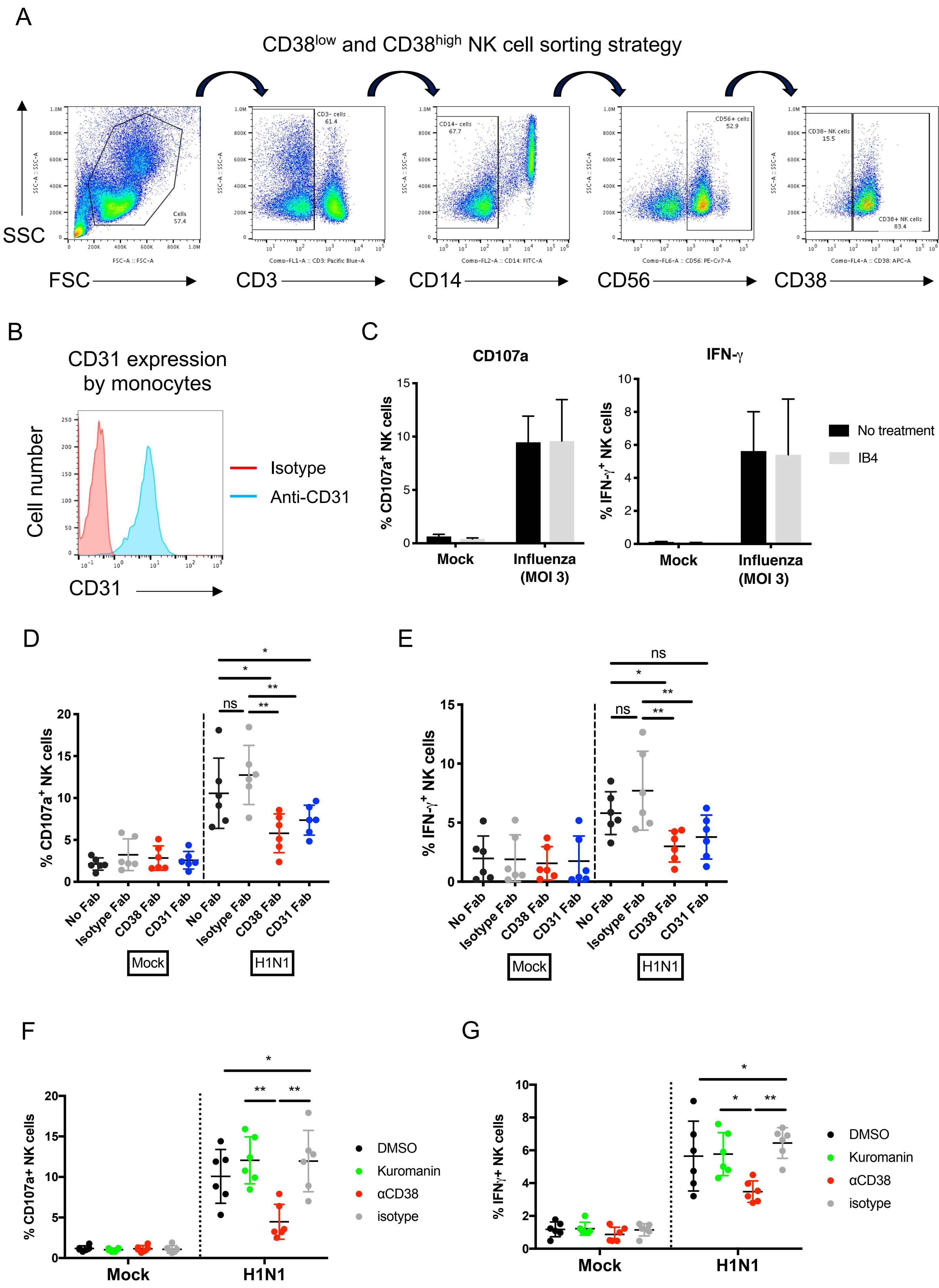

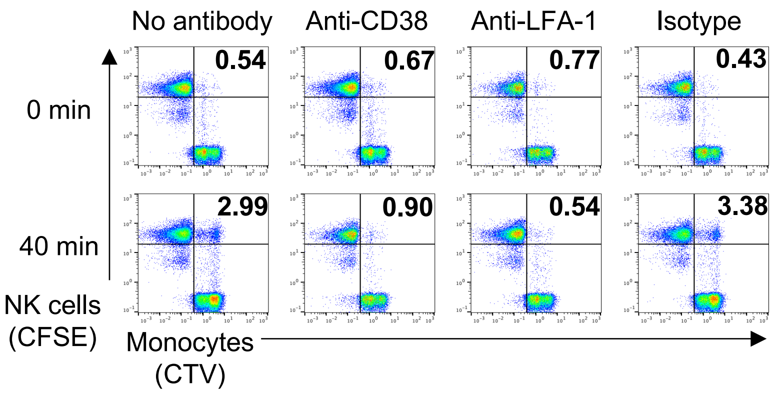

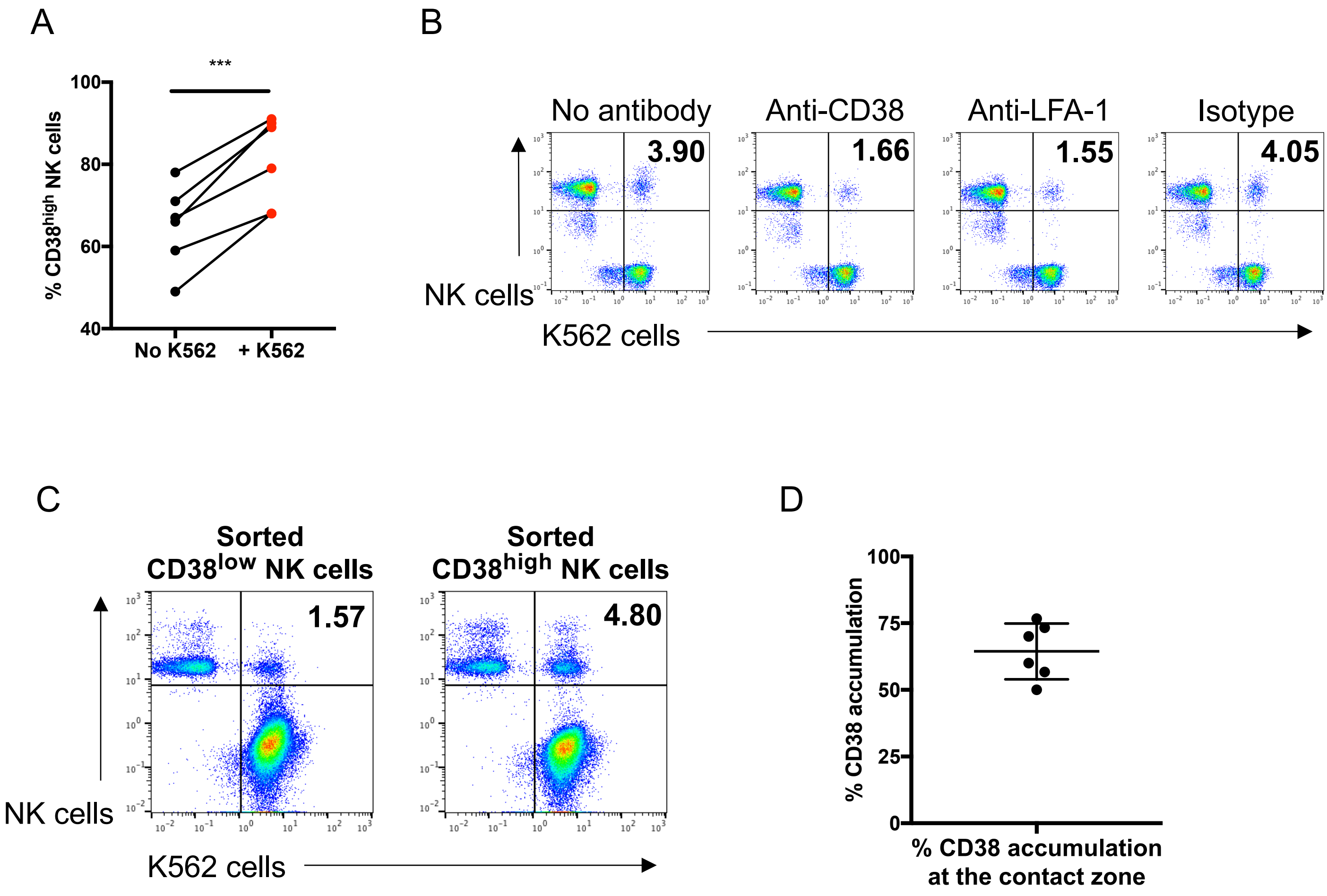
